# Clinically relevant SMAC mimetics do not enhance human T cell proliferation or cytokine production

**DOI:** 10.1101/2021.11.09.466489

**Authors:** Ashley Burton, Brittany Ligman, Claire Kearney, Susan E. Murray

**Affiliations:** Department of Biology, University of Portland, Portland, OR, United States; Department of Molecular Microbiology and Immunology, Oregon Health & Science University, Portland, OR, United States

## Abstract

Secondary mitochondria-derived activator of caspases (SMAC) mimetics are being tested in dozens of clinical trials to treat cancer. These targeted therapies mimic endogenous molecules that promote apoptosis by antagonizing inhibitors of apoptosis (IAPs), which are commonly overexpressed in cancer cells. In T cells, IAPs function to restrain non-canonical NF-kB signaling. Thus, it has been suggested that in addition to their direct anti-cancer mechanism of action, SMAC mimetics may activate T cells, thereby promoting anti-tumor immunity. Here, we tested the effect of three clinically relevant SMAC mimetics on the proliferation and activation of primary human T cells. As previously reported, SMAC mimetics killed tumor cells and activated non-canonical NF-kB in T cells at clinically relevant doses. Surprisingly, none of the SMAC mimetics augmented T cell proliferation or effector function. These results question the assumption that SMAC mimetics are likely to boost anti-tumor immunity in cancer patients.

## Introduction

In the past two decades, the number of targeted therapies for cancer treatment has grown exponentially. SMAC (secondary mitochondria-derived activator of caspases) mimetics comprise a class of targeted therapies that sensitize tumor cells to apoptosis^1,2^. They mimic an endogenous molecule, SMAC/Diablo, produced by mitochondria that antagonizes IAPs (inhibitor of apoptosis molecules), thus halting apoptosis downstream of intrinsic and extrinsic factors^3^. In non-cancerous cells, SMAC is released from the mitochondria and binds to IAPs, thereby allowing the cell to complete apoptosis via caspases. However, in cancer cells IAP upregulation is a tactic often used to evade apoptosis signals. Specifically, cellular IAPs (cIAPs) are essential to cancer cell survival as they bind directly to SMAC molecules and sequester them^4^. Furthermore, XIAPs are able to bind to caspases and halt apoptosis within the cancer cell completely.

The ability of SMAC mimetic to bind IAPs and therefore promote cancer cell apoptosis *in vitro* prompted interest in their clinical application^5–10^. In preclinical *in vivo* animal models, SMAC mimetics showed encouraging results, particularly when used in tandem with chemotherapy drugs to further sensitize cancer cells to chemotherapy^6,8–14^. However, in clinical trials, SMAC mimetics have yet to show efficacy as a monotherapy or to increase efficacy in combination with chemotherapy or immunotherapy^15^.

In addition to inhibiting apoptosis, IAPs also play a role in inhibiting non-canonical NF-kB activation in immune cells^16,17^. Thus, SMAC mimetics, which inhibit IAPs, would be predicted to increase non-canonical NF-kB activation. This is of particular relevance to cancer immunotherapy, because non-canonical NF-kB has been shown to be necessary for optimal T cell responses. In mouse models, activation of non-canonical NF-kB acts downstream of TNF receptor family members to promote T cell memory and effector functions^18–21^. Studies have shown that SMAC mimetics do, indeed, activate non-canonical NF-kB in immune cells^22–24^, and this is associated with enhanced T cell responses in mice *in vivo* and in humans *in vitro*^22,23^. In addition, SMAC mimetics have been shown to synergize with anti-PD-1 therapy in mouse models of cancer^25–27^, and this effect depended on the presence of T cells^26,27^. These studies suggest that SMAC mimetics might have two different anti-cancer mechanisms—a direct effect on cancer cell viability and an indirect effect via activating anti-tumor T cell responses. If true, this could have implications for combining SMAC mimetics with immunotherapy.

The aim of the present study was to directly compare the effect of several SMAC mimetics on multiple measures of human T cell function in CD4+ and CD8+ T cells. To our surprise, we found that none of the three SMAC mimetics tested affected T cell proliferation or CD25 expression under any culture conditions. Moreover, contrary to previous studies, none of the SMAC increased secreted or intracellular production of pro-inflammatory cytokines. Instead, birinapant, the SMAC mimetic farthest along in clinical trials, decreased percentages of multifunctional T cells that secrete two or more cytokines. These data suggest that, in humans, SMAC mimetics are unlikely to promote anti-tumor T cell responses, and thus may not synergize with immunotherapy as has been proposed.

## Methods

### Human Subjects

Peripheral blood mononuclear cell (PBMC) samples were from donor de-identified healthy adult males obtained via leukapheresis. Subjects were aged 22-56 and were seronegative for HIV, CMV, and Hepatitis B. The present study was determined by the University of Portland Institutional Review Board to be exempt under category #4, as we have no access to or ability to obtain identifying information about these donors.

### Antibodies and flow cytometry

The following antibodies were used for flow cytometry analysis: CD8 (BD Biosciences, HIT8a), CD4 (BD Biosciences, SK3), IFNγ (Invitrogen, 4S.B3), CD3 (eBioscience, SK7), TNFα (eBioscience, MAb11), IL-2 (Invitrogen, MQ1-17H12). The following antibodies were purchased from BioLegend: CD25 (BC96), CD3 (SK7), TNFα (MAb11), CD8 (RPA-T8), CD4 (OKT4). Stimulatory anti-CD3 (OKT3) and anti-CD28 (CD28.2) antibodies were from BioLegend. Flow cytometry was performed on a FACSymphony or LSR II (BD Biosciences). Data from flow cytometry was analyzed via FlowJo software (BD Biosciences).

### Media and other reagents

T cells were cultured in RPMI 1640 (Life Technologies) supplemented with 2 mM L-Glutamine (Life Technologies), 100 U/ml Penicillin (Invitrogen), 100 ug/ml Streptomycin (Invitrogen) and 10% FBS (R&D Systems). MDA-MB-231 breast cancer cells were grown in DMEM (Life Technologies) supplemented as above. FACS buffer consisted of 1x PBS, 2% FBS and 0.1% NaN3. Cell proliferation dye labeling buffer consisted of 1x PBS/0.1% (w/v) BSA (Sigma Aldrich). IAP antagonists (SMAC mimetics) birinapant, BV6, and LCL161 were from MedChem Express and Apex Bio, and were reconstituted at 10mM in DMSO and stored at –80° C.

### T Cell Culture

T cells were isolated from PBMCs with EasySep™ Human T cell Isolation Kit per manufacturer’s instructions (StemCell Technologies). Isolated T cells (0.5-1 × 10^5^ per well) or unseparated PBMCs (1 × 105 per well) were stimulated for 48-72 hours either with Dynabeads™ Human T Activator CD3/CD28 antibody coated beads (Thermo Fisher) at 2:1 or 4:1 bead:cells ratios or with plate-bound anti-CD3 and soluble anti-CD28, each at 2 ug/ml in the presence of varying concentrations of IAP antagonists. Cells were incubated at 37°C and 6.0% CO2. In some experiments, cells were labeled with CFSE or Cell Trace e450 proliferation dye (Thermofisher) prior to culture. Cells were incubated with 10uM CFSE or CTe450 in 1x PBS/0.1%BSA for 15 minutes at 37°C, then washed three times before culture.

### ICCS and Flow Cytometry

For proliferation measurements, cells were first stained with Zombie aqua fixable viability dye (Biolegend) for 15 minutes, then surface antibodies for 20 min, then washed and fixed in 1% paraformaldehyde before flow cytometric analysis. For apoptosis assessment of PBMC and T cells, following surface staining, cells were stained with Annexin V (R&D Systems) per manufacturer’s instructions and were run immediately without fixation or were fixed in 1% PFA diluted in Annexin V binding buffer, containing CaCl2 to preserve Annexin V binding. To measure intracellular cytokines, cells were treated with GolgiPlug™ (BD BioScience) for the last 5 hours of culture, then stained with fixable viability dye and surface antibodies as above. Cells were then permeabilized using Invitrogen eBioscience™ intracellular staining kit per manufacturer’s protocol and stained for IFNγ, TNFα and IL-2. Cells were run on a BD LSR II or BD Symphony and analyzed using FlowJo (BD Biosciences).

### Apoptosis of MDA-MB-231 cells

MDA-MB-231 cells were a kind gift from Dr. Pepper Schedin, OHSU. To assess apoptosis induced by IAP antagonists on these IAP overexpressing cells, MDA-MB-231 cells were cultured at in the presence of the indicated concentrations of IAP antagonist for 48 hours. Cells were assessed for apoptosis with Annexin V staining kit (R&D Systems) per manufacturer’s instructions and were analyzed immediately via flow cytometry.

### ELISA

IL-2 and IFNγ were assessed in cell culture supernatants via commercial ELISA. IL-2 Human Uncoated ELISA kit and IFNγ Human Uncoated ELISA kit (both from Invitrogen) and DuoSet Human IFN-gamma kit and Ancillary Reagent Kit 2 (R&D Systems) were performed according to manufacturer’s instructions. ELISA plates were read on iMark™ Microplate Reader (BioRad) and data analyzed with Excel.

### Western Blot

2 × 10^6^ T cells isolated from PBMC were stimulated with plate-bound anti-CD3 alone or anti-CD3 plus anti-CD28 for 24 or 48 hours. Cells were lysed in RIPA buffer and protein extracts from equal numbers of cells were resolved on a 12% SDS-PAGE gel (Biorad), followed by transfer onto Immun-Blot PVDF membranes (Biorad). Membranes were blocked in 5% non-fat dry milk and probed overnight with rabbit anti-NF-kB2 p100/p52 and GAPDH antibodies at 1:1000, followed by washing and 2 hour incubation with goat-anti-rabbit-HRP antibody at 1:5000 (all from Cell Signaling Technology). Membranes were washed again and detected with SuperSignal West Pico Chemiluminescent substrate (Thermofisher). Protein band intensities were quantified with ImageJ (NIH).

## Results

### SMAC Mimetics are able to activate the NF-kB pathway in human immune cells at concentrations that kill tumor cells

SMAC mimetics have been shown to antagonize IAPs, thereby activating the non-canonical NF-kB pathway^6,22–24^. To test the activation of NF-kB in human T cells, magnetically purified T cells were treated with birinapant and assessed via Western blot for the presence of p100/p52 isoforms. Increasing concentrations of birinapant activated non-canonical NF-kB in primary human T cells, as assessed by increased processing of NF-kB2 from the p100 to p52 isoform (figure 1A). In addition, birinapant exhibited no toxicity to T cells in doses up to at least 1uM (figure 1B). We believe this is physiologically relevant concentration, because in a clinical trial, birinapant concentrations in patient tissues averaged ~0.8 uM^28^. In contrast, SMAC mimetics in this study induced apoptosis of the SMAC-sensitive breast cancer cell line, MDA-MB-231 at >100-fold lower doses (figure 1C).

**Figure 1.**
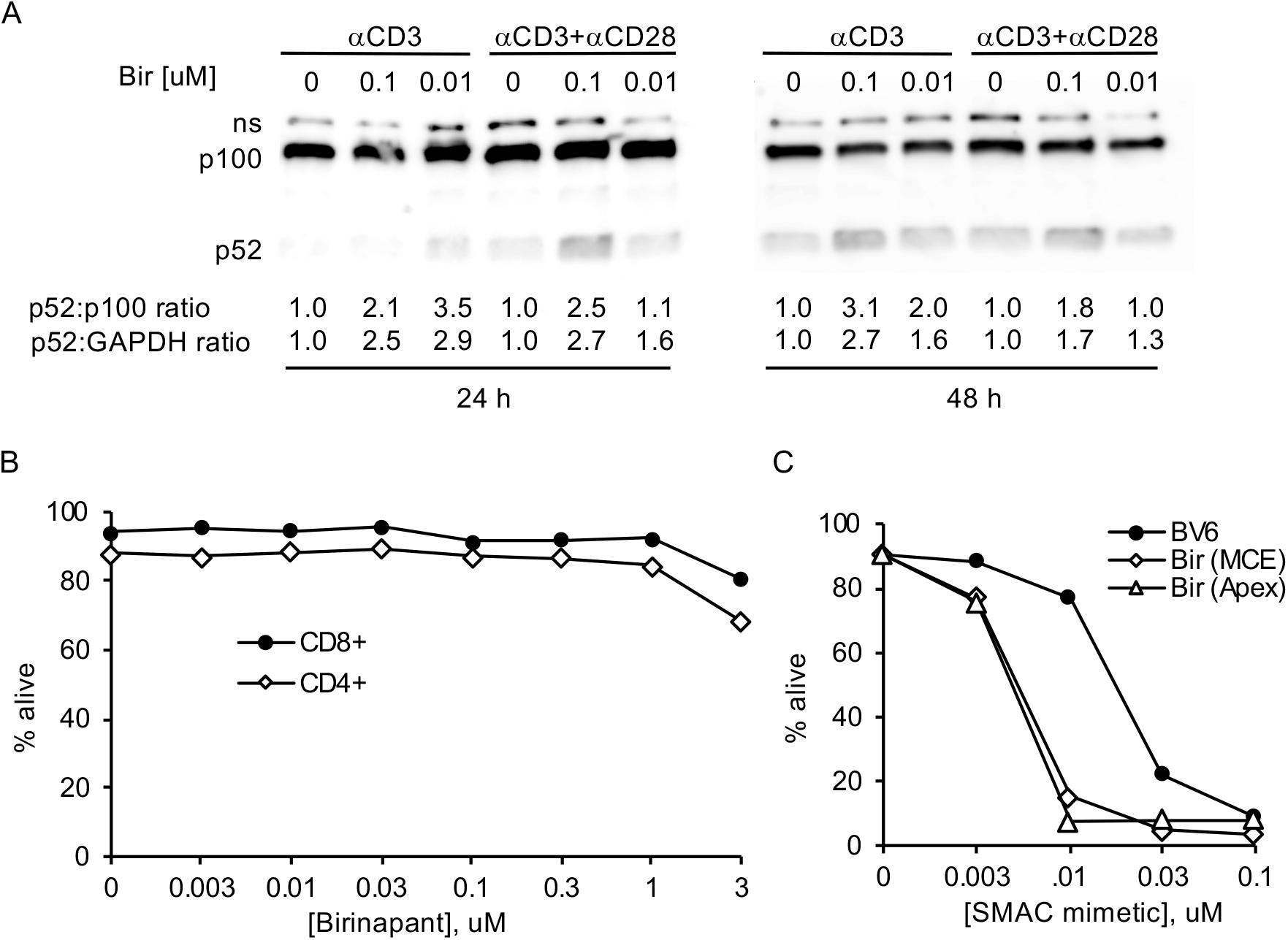
SMAC mimetics activate non-canonical NF-kB in human T cells at concentrations that kill cancer cells. A, T cells were magnetically purified from PBMC and stimulated with plate-bound anti-CD3 or anti-CD3/CD28 in the presence of the indicated concentrations of birinapant for 24 or 48 hours. Cells were harvested and total cellular protein was extracted, run on an SDS-PAGE gel, and blotted with an antibody to NF-kB2 that detects both inactive full-length (p100) and active cleaved (p52) isoforms. B, T cells were magnetically purified and stimulated with plate-bound anti-CD3/CD28 in the presence of the indicated concentrations of birinapant for 48 hours, then assessed for viability via Annexin/PI staining. C, MDA-MB231 breast cancer cell lines were cultured in the indicated concentrations of SMAC mimetic for 48 hours, then assessed for viability via Annexin/PI staining.

### SMAC mimetics do not increase T cell proliferation

Given that non-canonical NF-kB activation is intrinsically important for T cell clonal expansion^19,20^, we tested the effect of SMAC mimetics on T cell proliferation. In purified T cells, none of the three SMAC mimetics altered proliferation (figure 2B). Upon TCR stimulation, T cells upregulate the high affinity IL-2R, which is essential to allow them to robustly respond to autocrine IL-2. We assessed the effect of SMAC mimetics on CD25 expression in dividing T cells, and observed a trend towards decreased CD25 MFI with increasing doses of birinapant and LCL161 (figure 2C). This further suggests that SMAC mimetics do not enhance T cell proliferation. Since non-canonical NF-kB activation has also been shown to affect antigen presenting cells^29,30^, we considered that SMAC mimetics might affect T cell proliferation indirectly via effects on monocytes or dendritic cells. Thus, we tested the effect of SMAC mimetics on T cell proliferation in the context of total PBMC cultures. However, even in the presence of accessory cells, SMAC mimetics did not affect T cell proliferation (figure 2D) or CD25 expression (supplemental figure 1).

**Figure 2.**
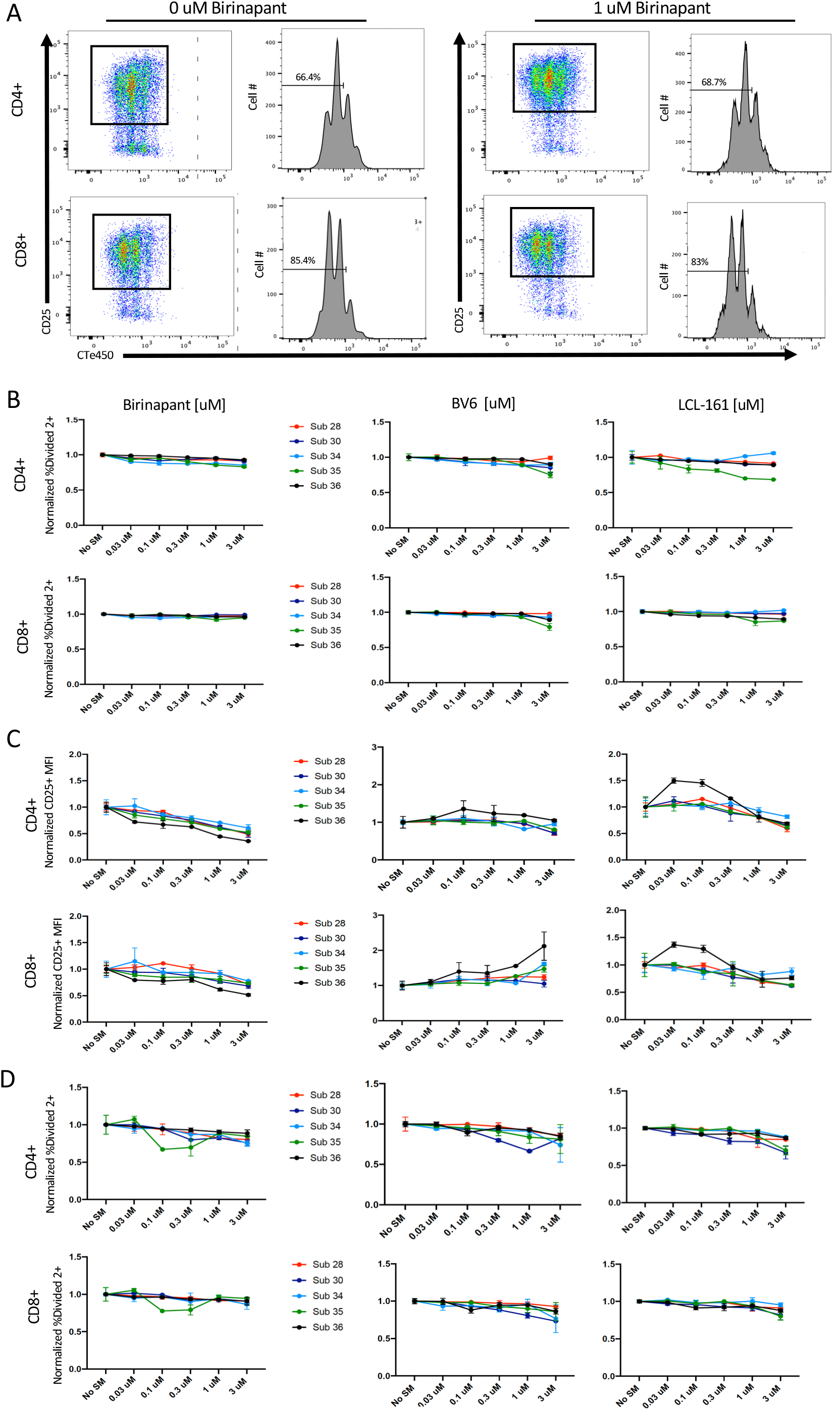
SMAC mimetics induce no change in T Cell or PBMC proliferation. **A,** Flow cytometry gating scheme. Cells were gated on CD25+ and CTe450 followed by CTe450 histograms for percent of cells divided. **B-C**, T cells were magnetically separated from whole PBMC samples, stained with CTe450 to detect cell division, and stimulated with Dynabeads at a concentration of 1:4 beads to cells in the presence of the indicated concentration of SMAC mimetic for 48 hours. Cells were then stained for surface antigens and analyzed via flow cytometry for percent of cells divided (**B**) and CD25 MFI (**C**). **D,** PBMCs were stained with CTe450 to detect cell division and stimulated with Dynabeads at a concentration of 1:4 beads to cells in the presence of the indicated SMAC mimetic at various concentration for 48 hours. Cells were then stained for surface antigens and analyzed via flow cytometry for percent of cells divided.

### SMAC mimetics do not increase secreted pro-inflammatory cytokine production in T cells

Previous studies showed that SMAC mimetics can increase IL-2 and TNFα production by T cells^22,23^. We tested the effect of SMAC mimetic on cytokine secretion from purified T cells and T cells stimulated in the context of PBMC. Culture supernatant was assessed for IL-2 and IFNγ by ELISA. Neither IL-2 nor IFNγ production was affected by birinapant in either culture condition (figure 3A). Likewise, cytokines were not increased in the presence of BV6 or LCL-161 (supplemental figure 2). Due to large subject to subject variation in cytokine concentrations, we also normalized each subjects’ cytokine production to the no SMAC mimetic control and plotted each subject’s response separately (figure 3B). These data show that SMAC mimetics had no effect on secreted IL-2 or IFNγ in stimulated T cells.

**Figure 3.**
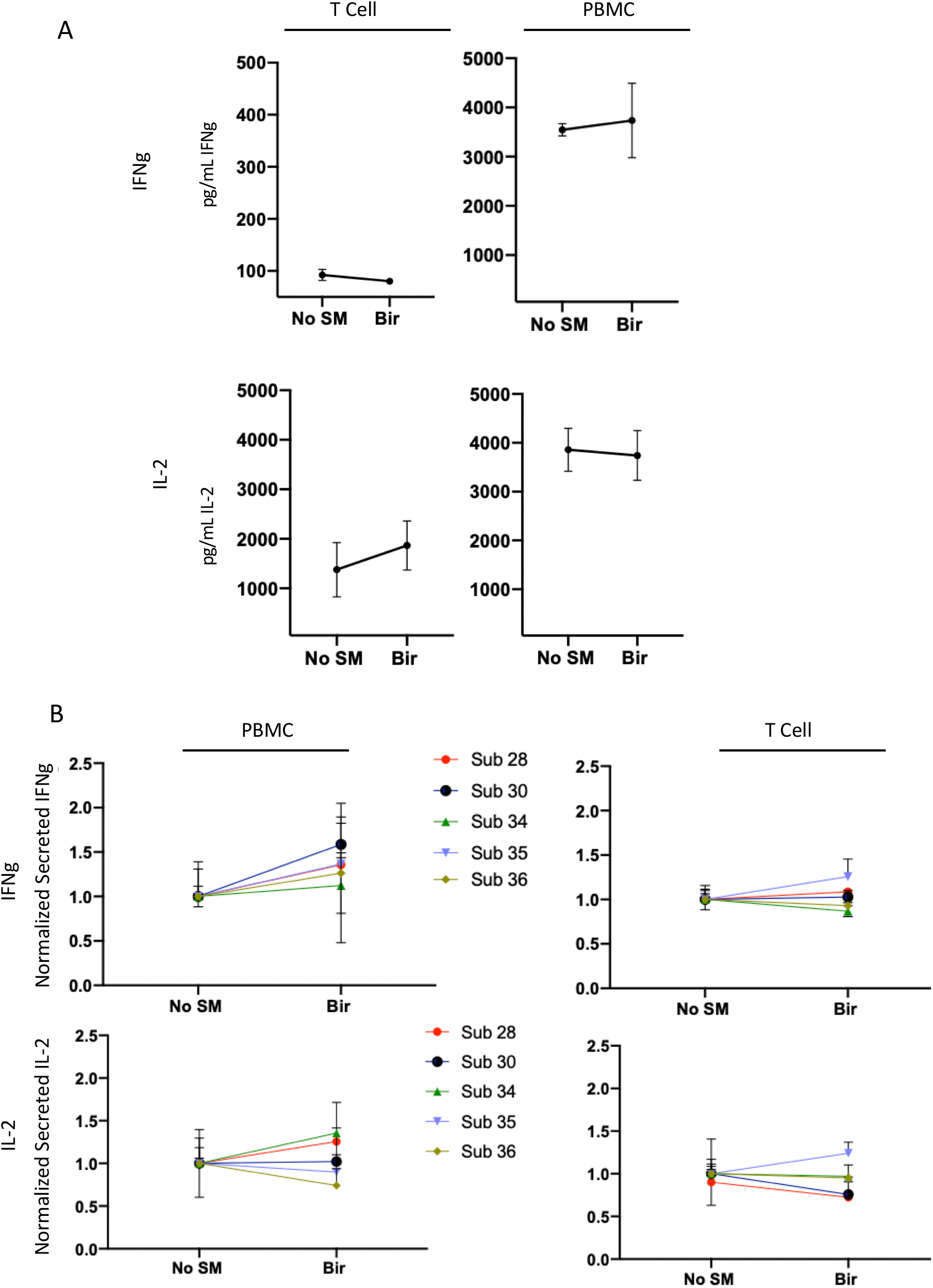
Birinapant does not affect IFNγ or IL-2 secretion. Unseparated PBMC or magnetically purified T cells were stimulated with Dynabeads at a 1:4 ratio beads to cells in the presence of 0 uM or 1 uM Birinapant. Supernatant was collected after 48 hours (IL-2) or 72 hours (IFNγ) and analyzed via ELISA. **A,** Average cytokine concentration in supernatants from 5 subjects +/− SD. **B,** Cytokine concentrations were normalized to each subject’s no SMAC mimetic control and plotted individually.

It was possible that SMAC mimetics altered cytokine production on a per cell basis, but that these changes were obscured by subsequent consumption of the cytokines produced. To directly assess cytokine production in individual CD4 and CD8 T cells, we performed ICCS on magnetically separated T cells as well as T cells cultured in the context of PBMCs. Birinapant did not alter the proportion of CD4 or CD8 T cells producing IL-2 or IFNγ (figure 4). To the contrary, birinapant actually decreased proportions of TNFα+/IFNγ+ double producers when CD4 or CD8 T cells were stimulated with plate-bound anti-CD3/CD28, although this trend did not hold true for bead-stimulated T cells (supplemental figure 3). Overall, we found no evidence that any of the SMAC mimetics we tested could increase cytokine production by human T cells.

**Figure 4.**
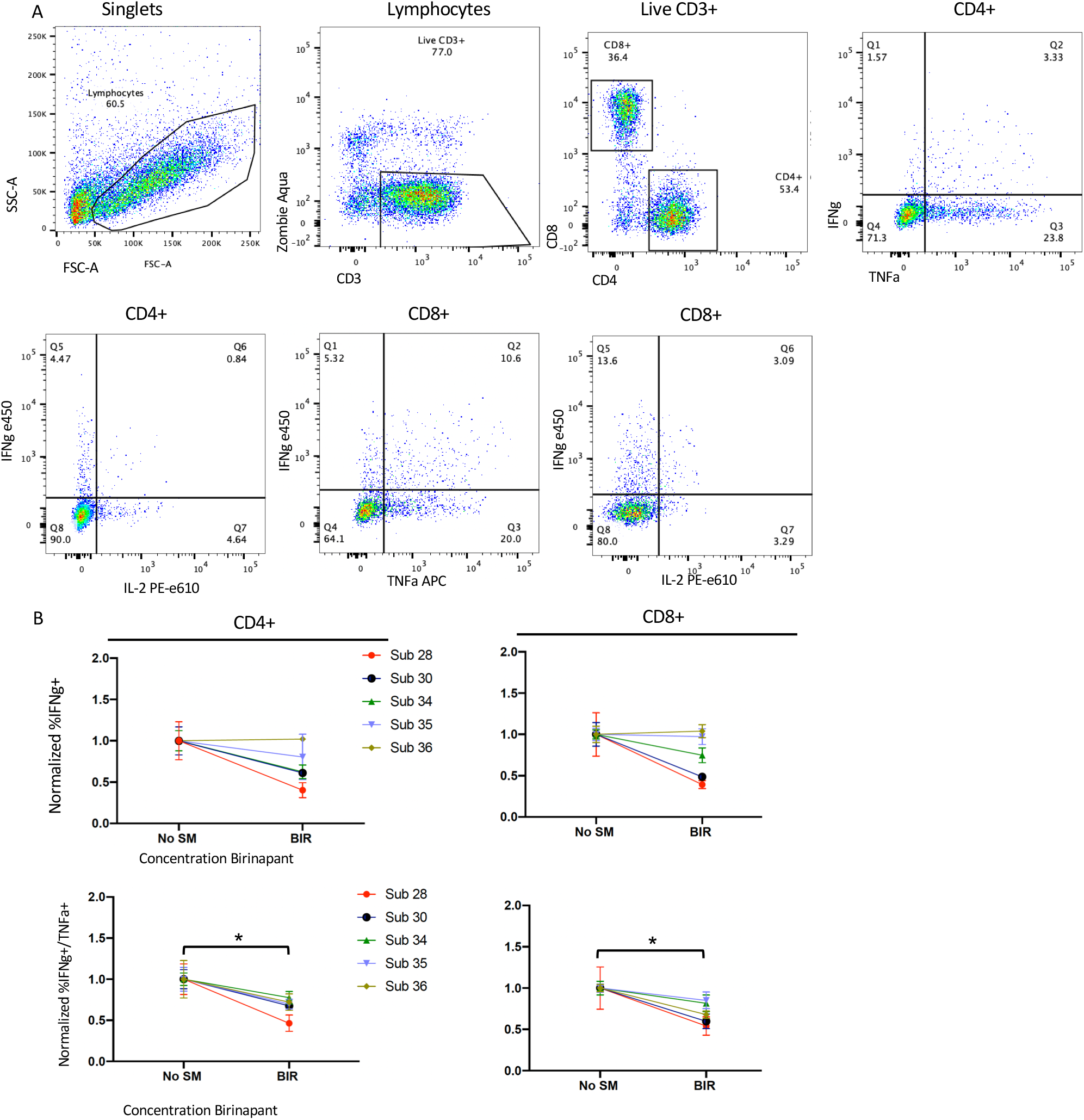
Birinapant does not affect the proportion of T cells producing cytokines. PBMC were stimulated with plate-bound anti-CD3/CD28 with or without 1 uM Birinapant for 48 hours, treated with BFA for 5 hours (without restimulation), stained for surface antigens and intracellular cytokines, and assessed via flow cytometry. **A,** Gating scheme for flow cytometry analysis. **B,** The percent of CD4 or CD8 T cells that were producing the indicated cytokine(s) was normalized to each subject’s no SMAC mimetic control and plotted individually. *Top*, total proportion of cells producing IFNγ. *Bottom*, proportion of cells producing both IFNγ and TNFα.

## Discussion

SMAC mimetics have been or are currently being tested in at least 34 clinical trials (ClinicalTrials.gov). In addition to their direct anti-cancer effects, it has been proposed that these drugs may augment T cell responses, thereby activating or boosting an anti-tumor immune response. Despite promising pre-clinical results in mouse models, to date SMAC mimetics have underperformed in clinical trials. Here, we sought to directly test the effect of several clinically relevant SMAC mimetics on human T cell function using multiple measures of T cell activation. Our data indicate that none of the drugs tested improved T cell responses in any of the numerous culture conditions and functional readouts we assessed. Instead, birinapant decreased proportions of CD4 and CD8 multi-functional cytokine producers, and both birinapant and LCL161 showed a trend towards decreasing CD25 expression by T cells, while none of the SMAC mimetics appreciably affected proliferation.

The SMAC mimetics we tested were biologically active as assessed by their ability to kill the SMAC mimetic-sensitive breast cancer cell line, MDA-MB-231, at very low concentrations. They were not toxic towards human T cells at clinically relevant concentrations up to 100 times higher than those that killed MDA-MB-231 cells. Furthermore, birinapant activated non-canonical NF-kB in human T cells as previously reported^22^. Thus, we can exclude the trivial explanations for our negative data that the SMAC mimetics we used were not biologically active or that toxic concentrations were used.

Previous studies have yielded contradictory data regarding the effect of SMAC mimetics on T cell proliferation. Dougan, *et al*, found that the SMAC mimetic, LBW-242, modestly increased mouse T cell proliferation *in vitro*^22^; whereas, Gentle, *et al*, observed that LBW-242 had the opposite effect on mouse T cell proliferation *in vitro* and *in vivo* in the context of a viral infection^31^. LCL161 enhanced *in vitro* expansion of human T cells specific for some antigens, but not others^23^. Moreover, the latter study reported that LCL161 did not alter proliferation of human T cells in response to polyclonal stimulation, a result that mirrors our own. Likewise, the effects of SMAC mimetics on cytokine production by T cells are somewhat discordant. LBW-242 and LCL161 increased IL-2 production by TCR stimulated mouse and human T cells, respectively, and LCL161 also increased TNF□ production by human T cells^22,23^. However, SMAC mimetics may have divergent effects on different helper T cell subsets. Rizk, *et al*, found that SMAC mimetics decreased IL-17 production (Th17), increased IL-9 and IL-13 production (Th9 and Th2), and did not alter IFN□ or TNF□ production (Th1) in mouse T cells^24^. This is highly relevant to cancer therapy, as type 1 responses from both Th1 and CD8 T cells are considered to provide the most robust anti-tumor immunity. We therefore focused on type 1 cytokines— IL-2, IFNγ, and TNFα—and found no evidence that the SMAC mimetics we tested increased production of any of these cytokines. Importantly, we tested multiple SMAC mimetics across a wide range of clinically relevant drug concentrations under several different stimulation conditions, and we measured both intracellular and secreted cytokines. The only significant effect on T cell cytokine production we observed was a decrease in multi-functional T cells that could secrete more than one cytokine.

Given that we observed increased non-canonical NF-kB activation upon SMAC mimetic treatment of human T cells, it is interesting that this did not lead to enhanced proliferation or cytokine production. One possibility is that non-canonical NF-kB simply wasn’t activated sufficiently to boost T cell responses. Another is that non-canonical NF-kB activation may play a lesser role in activation of human versus mouse T cells. This is supported by the phenotype of patients with biallelic loss of function mutations in NIK or IKKα, both central kinases in the non-canonical NF-kB pathway^32,33^. These patients had severe B cell defects and lack of secondary lymphoid structures, but no alterations in T cell subsets or in T cell proliferation in response to anti-CD3 or mitogen stimulation. Although they had reduced T cell recall responses to TT and PPD antigens, this was likely the result of poor T cell priming due to the absence of lymph nodes in these patients. Finally, SMAC mimetics may have targets other than xIAP, cIAP1, and cIAP2 in T cells. In cancer cells, LCL161 has been shown to directly modulate ABCB1/MDR1-ATPase and decrease intracellular ATP levels^34^. Proliferating and activated T cells are highly dependent on intracellular ATP^35^; thus if SMAC mimetics decrease ATP in human T cells, this effect could counteract potential activating effects of SMAC mimetic-induced non-canonical NF-kB.

Our data challenge the assumption that SMAC mimetics are likely to boost human T cell responses to improve tumor immunity. They also suggest that combining SMAC mimetics with checkpoint blockade or other immunotherapy may be additive, rather than synergistic. This is relevant to the design of future clinical trials as SMAC mimetics continue to advance through the pipeline of targeted cancer therapies.

## Author Contributions

S.E.M. conceived of the project, planned the experiments, performed and analyzed experiments, and wrote the manuscript. A.B. performed and analyzed experiments, created figures, and wrote the manuscript. B.L. and C.K performed and analyzed experiments.

## Acknowledgements

We would like to thank Ann B. Hill for providing PBMC, Pepper Schedin for MDA-MB-231 cells, and Michael Munks for helpful discussions. Funding was provided by the M.J. Murdock Charitable Trust and Medical Research Foundation of Oregon.

## Notes

### Competing Interest Statement

The authors have declared no competing interest.

## References

1. Morrish, E., Brumatti, G. & Silke, J. Future Therapeutic Directions for Smac-Mimetics. Cells 9, (2020).

2. Fulda, S. & Vucic, D. Targeting IAP proteins for therapeutic intervention in cancer. Nat Rev Drug Discov 11, 109–124 (2012).

3. Du, C., Fang, M., Li, Y., Li, L. & Wang, X. Smac, a mitochondrial protein that promotes cytochrome c-dependent caspase activation by eliminating IAP inhibition. Cell 102, 33–42 (2000).

4. Gyrd-Hansen, M. & Meier, P. IAPs: from caspase inhibitors to modulators of NF-kappaB, inflammation and cancer. Nat Rev Cancer 10, 561–574 (2010).

5. Benetatos, C. A. et al. Birinapant (TL32711), a bivalent SMAC mimetic, targets TRAF2-associated cIAPs, abrogates TNF-induced NF-κB activation, and is active in patient-derived xenograft models. Mol Cancer Ther 13, 867–879 (2014).

6. Carter, B. Z. et al. Synergistic targeting of AML stem/progenitor cells with IAP antagonist birinapant and demethylating agents. J Natl Cancer Inst 106, djt440 (2014).

7. Li, L. et al. A small molecule Smac mimic potentiates TRAIL- and TNFalpha-mediated cell death. Science 305, 1471–1474 (2004).

8. Probst, B. L. et al. Smac mimetics increase cancer cell response to chemotherapeutics in a TNF-α-dependent manner. Cell Death Differ 17, 1645–1654 (2010).

9. Greer, R. M. et al. SMAC mimetic (JP1201) sensitizes non-small cell lung cancers to multiple chemotherapy agents in an IAP-dependent but TNF-α-independent manner. Cancer Res 71, 7640–7648 (2011).

10. Krepler, C. et al. The novel SMAC mimetic birinapant exhibits potent activity against human melanoma cells. Clin Cancer Res 19, 1784–1794 (2013).

11. Janzen, D. M. et al. An apoptosis-enhancing drug overcomes platinum resistance in a tumour-initiating subpopulation of ovarian cancer. Nat Commun 6, 7956 (2015).

12. Dineen, S. P. et al. Smac mimetic increases chemotherapy response and improves survival in mice with pancreatic cancer. Cancer Res 70, 2852–2861 (2010).

13. Lalaoui, N. et al. Targeting triple-negative breast cancers with the Smac-mimetic birinapant. Cell Death Differ 27, 2768–2780 (2020).

14. Lu, J. et al. Therapeutic potential and molecular mechanism of a novel, potent, nonpeptide, Smac mimetic SM-164 in combination with TRAIL for cancer treatment. Mol Cancer Ther 10, 902–914 (2011).

15. Chang, Y. C. & Cheung, C. H. A. An updated review of SMAC mimetics, LCL161, Birinapant, and GDC-0152 in cancer treatment. Applied Sciences 11, 335 (2021).

16. Vallabhapurapu, S. et al. Nonredundant and complementary functions of TRAF2 and TRAF3 in a ubiquitination cascade that activates NIK-dependent alternative NF-kappaB signaling. Nat Immunol 9, 1364–1370 (2008).

17. Zarnegar, B. J. et al. Noncanonical NF-kappaB activation requires coordinated assembly of a regulatory complex of the adaptors cIAP1, cIAP2, TRAF2 and TRAF3 and the kinase NIK. Nat Immunol 9, 1371–1378 (2008).

18. Aya, K. et al. NF-(kappa)B-inducing kinase controls lymphocyte and osteoclast activities in inflammatory arthritis. J Clin Invest 115, 1848–1854 (2005).

19. Rowe, A. M. et al. A cell-intrinsic requirement for NF-kappaB-inducing kinase in CD4 and CD8 T cell memory. J Immunol 191, 3663–3672 (2013).

20. Murray, S. E. et al. NF-kappaB-inducing kinase plays an essential T cell-intrinsic role in graft-versus-host disease and lethal autoimmunity in mice. J Clin Invest 121, 4775–4786 (2011).

21. Jin, W., Zhou, X. F., Yu, J., Cheng, X. & Sun, S. C. Regulation of Th17 cell differentiation and EAE induction by MAP3K NIK. Blood 113, 6603–6610 (2009).

22. Dougan, M. et al. IAP inhibitors enhance co-stimulation to promote tumor immunity. J Exp Med 207, 2195–2206 (2010).

23. Knights, A. J., Fucikova, J., Pasam, A., Koernig, S. & Cebon, J. Inhibitor of apoptosis protein (IAP) antagonists demonstrate divergent immunomodulatory properties in human immune subsets with implications for combination therapy. Cancer Immunol Immunother 62, 321–335 (2013).

24. Rizk, J. et al. SMAC mimetics promote NIK-dependent inhibition of CD4+ Th17 cell differentiation. Sci Signal 12, (2019).

25. Kearney, C. J. et al. PD-L1 and IAPs co-operate to protect tumors from cytotoxic lymphocyte-derived TNF. Cell Death Differ 24, 1705–1716 (2017).

26. Chesi, M. et al. IAP antagonists induce anti-tumor immunity in multiple myeloma. Nat Med 22, 1411–1420 (2016).

27. Beug, S. T. et al. Smac mimetics synergize with immune checkpoint inhibitors to promote tumour immunity against glioblastoma. Nat Commun 8, (2017).

28. Noonan, A. M. et al. Pharmacodynamic markers and clinical results from the phase 2 study of the SMAC mimetic birinapant in women with relapsed platinum-resistant or -refractory epithelial ovarian cancer. Cancer 122, 588–597 (2016).

29. Lind, E. F. et al. Dendritic cells require the NF-kappaB2 pathway for cross-presentation of soluble antigens. J Immunol 181, 354–363 (2008).

30. Hofmann, J., Mair, F., Greter, M., Schmidt-Supprian, M. & Becher, B. NIK signaling in dendritic cells but not in T cells is required for the development of effector T cells and cell-mediated immune responses. J Exp Med 208, 1917–1929 (2011).

31. Gentle, I. E. et al. Inhibitors of apoptosis proteins (IAPs) are required for effective T-cell expansion/survival during antiviral immunity in mice. Blood 123, 659–668 (2014).

32. Willmann, K. L. et al. Biallelic loss-of-function mutation in NIK causes a primary immunodeficiency with multifaceted aberrant lymphoid immunity. Nat Commun 5, 5360 (2014).

33. Bainter, W. et al. Combined immunodeficiency with autoimmunity caused by a homozygous missense mutation in inhibitor of nuclear factor κB kinase alpha. Sci Immunol 6, 6723 (2021).

34. Chang, Y. C. et al. The SMAC mimetic LCL161 is a direct ABCB1/MDR1-ATPase activity modulator and BIRC5/Survivin expression down-regulator in cancer cells. Toxicol Appl Pharmacol 401, 115080 (2020).

35. Shyer, J. A., Flavell, R. A. & Bailis, W. Metabolic signaling in T cells. Cell Res 30, 649–659 (2020).

