## Supplemental figures 1-3 for "Clinically relevant SMAC mimetics do not enhance human T cell proliferation or cytokine production"

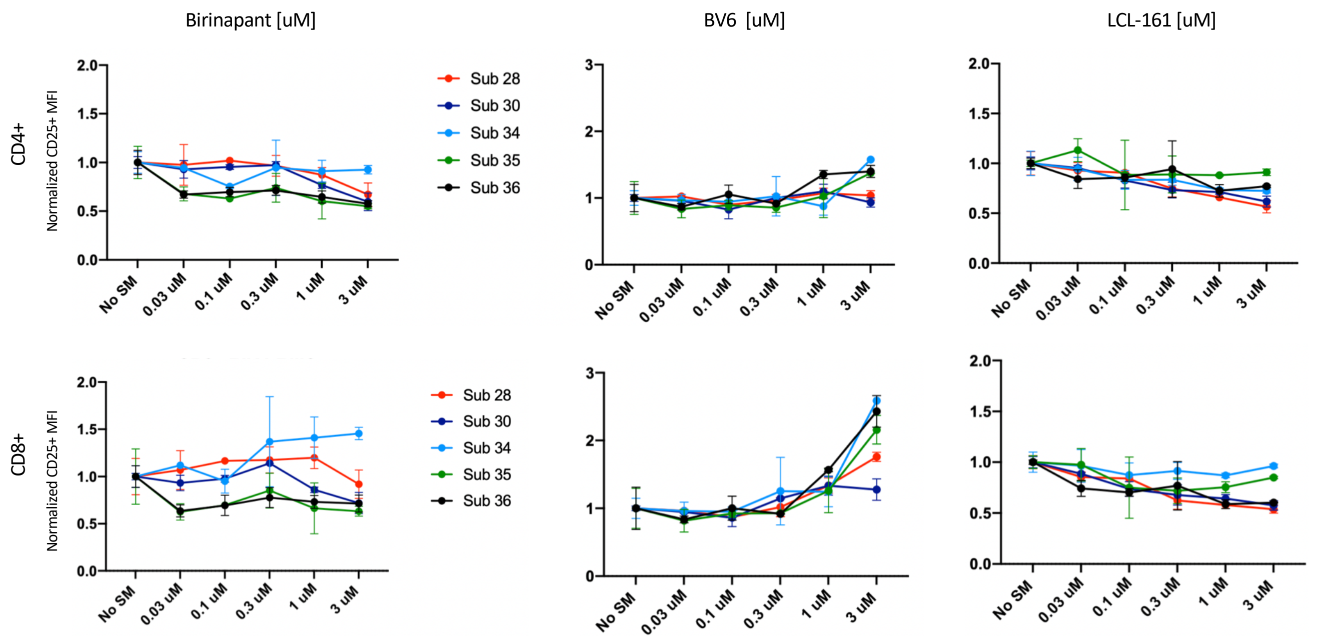


Supplementary figure 1. SMAC mimetics do not affect CD25 expression by T cells cultured with PBMC. PBMCs were stimulated with Dynabeads at a concentration of 1:4 beads to cells in the presence of the indicated SMAC mimetic for 48 hours. Cells were stained for surface markers and analyzed via flow cytometry for CD25 expression.

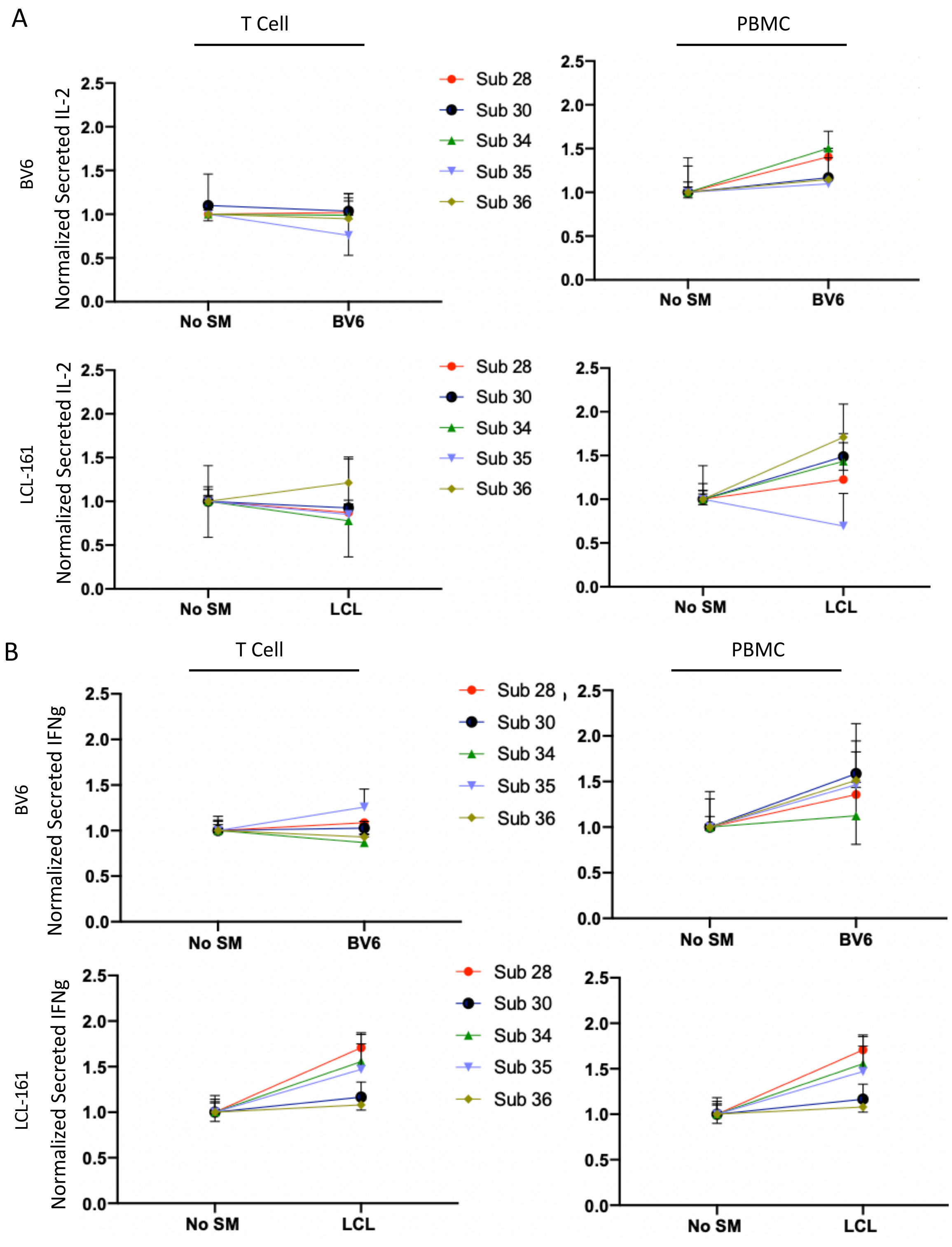


Supplemental figure 2. BV6 and LCL-161 do not affect secreted IFNγ or IL-2. Magnetically purified T cells or total PBMC were stimulated with Dynabeads at a 1:4 ratio beads to cells in the presence or absence of the indicated SMAC mimetic. Supernatant was collected after 48 or 72 hours for IL-2 (**A**) or IFNγ (**B**) assessment, respectively. Supernatants were analyzed via ELISA and normalized to each subject's no SMAC mimetic.

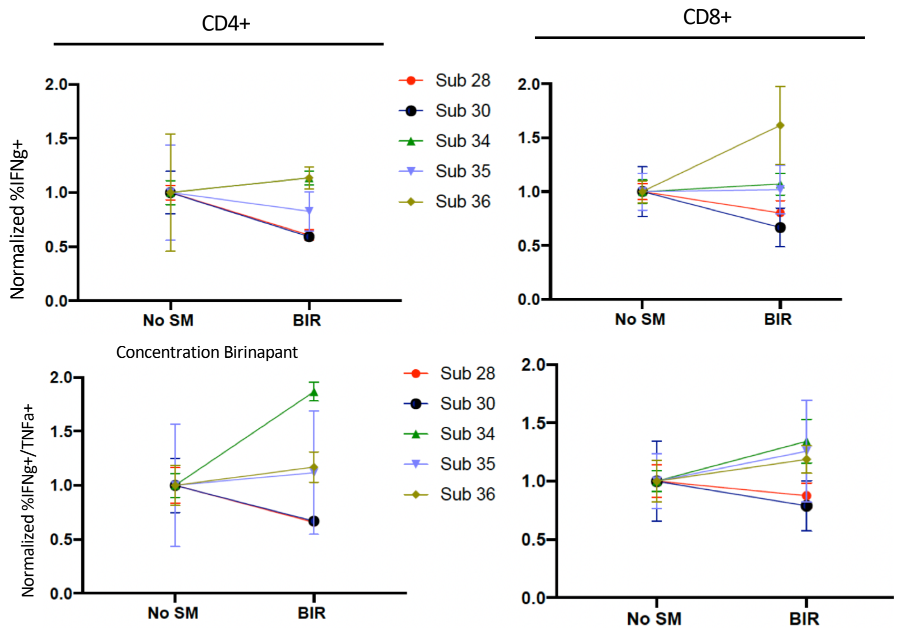


Supplemental figure 3. Birinapant does not affect intracellular cytokine production in bead stimulated PBMC samples. Unseparated PBMCs were stimulated with a 1:4 ratio of Dynabeads:cells in the absence or presence of Birinapant for 48 hours. Cells were then treated with BFA for 5 hours (without further stimulation), stained for surface antigens and intracellular cytokines and analyzed by flow cytometry. Percent of CD4 or CD8 T cells that were positive for the indicated cytokine(s) were normalized to each subject's no SMAC mimetic.
